# Proteome analysis reveals common players between the physiological neurodegeneration of the ascidian *Ciona intestinalis* and the pathological neurodegeneration in humans

**DOI:** 10.64898/2026.01.12.698666

**Authors:** Daniele Capitanio, Silvia Mercurio, Ettore Mosca, Cristina Battaglia, Marco Venturin, Roberta Pennati

**Author notes:** Corresponding author Department of Environmental Science and Policy Università degli Studi di Milano. these authors equally contributed to the work (Marco Venturin, Roberta Pennati).

## Abstract

Tunicates, including ascidians, are recognized as the true ‘sister group’ of vertebrates and are emerging as models to study the development and degeneration of central nervous system (CNS). Ascidian larvae have the typical chordate body plan that includes a dorsal neural tube. During their metamorphosis, a deep tissue reorganization takes place, with some tissues that degenerate while others develop to become functional during the adult life. The larval CNS also degenerates and most neurons disappear, making room for the formation of adult CNS. The genome of the ascidian *Ciona intestinalis* has been sequenced and annotated, with several CNS specific genes that have been characterized, revealing specification mechanisms shared with humans. These features make ascidian metamorphosis a good model to study the mechanisms underlying physiological CNS degeneration and to compare them to the pathological conditions typical of neurodegenerative diseases.

In order to shed light on the molecular determinants of *C. intestinalis* metamorphosis and neurodegeneration, we analyzed the proteome at three stages of development: swimming larva (SwL, Hotta stage 28), settled larva (SetL, Hotta stage 32) and metamorphosing larva (MetL, Hotta stage 34). A total of 405 modulated proteins were identified by mass spectrometry by comparing the three stages. Enrichment and network analysis showed the involvement of several processes/pathways, including autophagy and mTOR pathway, and actin cytoskeleton organization and remodeling among the most significant ones.

This study elucidates molecular pathways underlying ascidian metamorphosis and highlights shared mechanisms between physiological neurodegeneration in ascidians and pathological neurodegeneration in humans.

## INTRODUCTION

Neurodegenerative diseases are among the main causes of an increasing proportion of morbidity and mortality in developed countries. Due to an increased life expectancy, pathologies such as Alzheimer’s disease (AD) and Parkinson disease (PD) are becoming more common. Even though these syndromes appear different in many aspects, there is increasing evidence that they share some key pathological mechanisms (Skovronsky, Lee, and Trojanowski 2006).

Several model systems have been employed to study neurodegeneration processes (Götz et al. 2004; Link 2005). Human cell lines constitute a precious system to study specific mechanisms associated with these pathologies, but cells cannot reproduce the complex interactions among cell types present in an organism (Cetin et al. 2022). For this reason, researchers employed a variety of animal models, assuming that the basic processes of human neurodegeneration were present in these organisms. Transgenic murine models made an important contribution to this research field (Zhong et al. 2024) but there are several limits for their use, including ethical issues and the cost and time for their production. Thus, an increasing number of researchers took advantage in the use of invertebrate models, mainly *Drosophila* (Prüßing, Voigt, and Schulz 2013) and *C. elegans* (Alvarez et al. 2022) for their fast development and their highly characterized genomes. Besides these advantages, the long evolutionary distance from vertebrates makes it difficult to extrapolate results obtained with these models to human diseases.

Tunicates have been recognized as the sister group of vertebrates with which they share an exclusive common ancestor (Delsuc et al. 2006) and are emerging as excellent models to study the development, aging and degeneration of central nervous system (CNS) (Jeffery 2015; Virata and Zeller 2010). Ascidians are the most numerous group of tunicates; they are sessile filter feeder animals that develop through a swimming lecithotrophic tadpole larva. The larva retains the main key characters of the chordate body plan including a dorsal hollow neural tube that lays dorsal to the notochord (Passamaneck and Di Gregorio 2005). The larval CNS of *Ciona intestinalis* (here after Ciona), one of the most studied ascidians, consists of less than 100 neurons and 250 glial cells (Nicol and Meinertzhagen 1991). It contains suitably few cells to enable the final counts, their identification, the developmental history and the analysis of functions cell by cell (Meinertzhagen 2005). The CNS can be subdivided morphologically and functionally along the anteroposterior axis into four regions similarly to the vertebrate one: the sensory vesicle with the sensory organs, the neck, the visceral ganglion and the tail nerve cord. Beside its small size, it includes several neuronal types, distinguishable by the neurotransmitter that they utilize (Manni and Pennati 2016). Moreover, the CNS of ascidians develops through a process of neurulation that takes place at the end of the gastrula period and starts with the differentiation of a neural plate (Nishida and Stach 2014).

After a short swimming period, the ascidian larva undergoes a deep morphological change and becomes sessile (Hotta, Dauga, and Manni 2020). The metamorphic process is characterized by a rearrangement of larval tissue and organs, some of which degenerate and are reabsorbed whereas others develop and become functional for adult life. The CNS degenerates meanwhile the adult CNS differentiates (Hotta, Dauga, and Manni 2020).

The functions of larval and adult CNS are completely different, since in the larva stage CNS mainly coordinates swimming behavior whereas it orchestrates a complex of organs in the sessile adults. The CNS of adult ascidians consists of a ganglion located between the two siphons. Five main nerves branch off the ganglion: a pair of anterior nerves, a pair of posterior nerves, and a single posterior-ventral visceral nerve. During metamorphosis, the larval CNS is deeply remodeled with extensive loss of neurons. This loss occurs during the first stages of the metamorphosis, at the beginning of the body axis rotation stage (Hotta, Dauga, and Manni 2020). Cell-type-specific tracing with the fluorescent protein Kaede showed that most of the larval neurons disappear during metamorphosis. The cholinergic neurons of the motor ganglion are exceptions: a few cholinergic neurons remain after metamorphosis and are integrated into the adult CNS (Horie et al. 2011). New adult neurons differentiate from the neurohypophysial duct, a structure connecting the CNS to the oral siphon that is outside to the main part of the CNS, where cells of the wall undergo extensive mitosis (Dufour et al. 2006). Numerous apoptotic cells are detected in the larval CNS during metamorphosis (Tarallo and Sordino 2004), suggesting that the apoptosis pathway plays a role in the process of larval neurons loss.

The genome of Ciona was published in 2002 (Dehal et al. 2002). It is considered quite simple since the size is approximately 160 mega base pairs (Mbp) per haploid and contains approximately 16.000 genes. Since the genome was published, its wide analyses have successfully characterized an increasing number of genes specifically expressed in the CNS revealing molecular pathway shared with humans (Mochizuki, Satou, and Satoh 2003). Virata and Zeller enlightened the potentialities of Ciona as a useful model to study AD. Indeed, its genome retains orthologues of all the genes involved in the processing of Amyloid Precursor Protein (APP) including *BACE1* that is not present in other invertebrate models. Moreover, transgenic larvae can process human APP, but transgenic ones expressing a mutated Aβ in the nervous system were unable to attach to the substrate and to metamorphose (Virata and Zeller 2010).

These characteristics make the ascidian metamorphosis an exceptional model to study the mechanisms that regulate neurodegeneration in physiological conditions and to compare them with pathological conditions in neurodegenerative diseases.

In this paper, we compared the proteomes of swimming larvae, individuals at the onset of metamorphosis, and newly metamorphosed juveniles to identify differentially expressed proteins involved in physiological neurodegeneration. Our results identified proteins exhibiting trends comparable to those observed in human neurodegenerative disorders, revealing a remarkable similarity between physiological neurodegeneration in ascidians and pathological neurodegeneration in humans.

## MATERIALS AND METHODS

### Animals

Adults of *Ciona intestinalis* were collected by the fishery service of the Station Biologique de Roscoff (France). Once in the laboratory, animals were maintained in aquaria at 18°C for two days for acclimatization. To obtain synchronously developing embryos, in vitro fertilization was performed with gametes collected by dissection from five adults. Embryos were reared at 18°C in artificial seawater supplemented with 0.1 mM HEPES (Sigma, Italy) (AFWH) until they reached the larva stage. Since after hatching the development is no more synchronous, hatched larvae were carefully monitored by a stereo microscope to identify individuals that reached the selected developmental stages. When samples reached the desired developmental stage, they were manually collected, pooled in groups of 200 individuals, and immediately frozen in liquid nitrogen. The fertilization was replicated three times to obtain three biological replicas.

To catch the molecular events of neurodegeneration, we selected three developmental stages: i) swimming larva stage (SwL, stage 28 of Hotta) (Hotta, Dauga, and Manni 2020), characterized by elongated papillae and a square shaped trunk. The larvae were actively swimming with vigorous tail movements. This stage was reached about 2 hours post hatching; ii) settled larva (SetL corresponding to Mid Tail Absorption Stage 32 of Hotta), the larva was attached to the substrate by means of the adhesive papillae. The tail was not moving and at least 50% of the tail was resorbed into the larval trunk. This stage was reached 10-11 hours after hatching; iii) metamorphosed larva (MetL, corresponding to the Early Body Axis Rotation Stage 34 of Hotta), the stalk was elongated and formed an angle of about 90° with the endostyle axis. This stage was reached 14-18 hours after hatching.

### Protein extraction and proteomic analysis

A total of 9 samples were collected from three stages of larval development of Ciona (3 biological replicates for each stage) and their proteins were extracted in lysis buffer (0.1 M Tris/HCl pH 7.6, 4% SDS, 0.1 M DTT and 0.001 M PMSF) through sonication. Proteins were denatured at 95°C for 3 min and lysates clarified by centrifugation at 16000 g for 15 min. Protein concentrations were determined using the 2D-Quant Kit (Cytiva) and 120 µg of each extract were processed following the Filter aided sample preparation (FASP) method (Wiśniewski et al. 2009). Then liquid chromatography coupled to electrospray tandem mass spectrometry (LC–ESI–MS/MS) with label-free quantification analysis was performed. All samples were analyzed in triplicate at UNITECH OMICs (University of Milano, Italy) using a Dionex Ultimate 3000 nano-LC system (Sunnyvale CA, USA) connected to an Orbitrap Fusion™ Tribrid™ Mass Spectrometer (Thermo Scientific, Bremen, Germany) equipped with nano electrospray ion source. Peptide mixtures were pre-concentrated onto an Acclaim PepMap 100 – 100 μm x 2 cm C18 (Thermo Scientific) and separated on EASY-Spray column ES802A, 25 cm x 75 μm ID packed with Thermo Scientific Acclaim PepMap RSLC C18, 3 μm, 100 Å using mobile phase A (0,1 % formic acid in water) and mobile phase B (0.1% formic acid in acetonitrile 20/80, v/v) at a flow rate of 0.300 μL/min. The temperature was set to 35°C. The sample injection volume was 3 μL. MS spectra were collected over an m/z range of 375 – 1500 Da at 120,000 resolutions, operating in the data dependent mode (TOP 40). HCD was performed with collision energy set at 35 eV. Polarity: positive.

### Bioinformatic and statistical analysis

#### Protein annotation and differential expression analysis

Proteins were identified to match with *Ciona intestinalis* UNIPROT reference proteome (UP000008144, https://www.uniprot.org/proteomes/UP000008144) and their peaks were quantified with MaxQuant software (version 2.7.0.0, Max Planck Institute of Biochemistry, Germany) (Cox and Mann 2008). Quantification in MaxQuant was performed using the built-in extracted ion chromatogram (XIC)-based label-free quantification (LFQ) algorithm using fast LFQ (Cox et al. 2014). Detection of differentially expressed proteins (DEPs) was performed with Perseus software (Tyanova et al. 2016) (version 2.0.11, Max Planck Institute of Biochemistry). ANOVA and Tukey post-hoc test (p-value < 0.05) were used to obtain DE proteins with their corresponding log fold change (LFC) values for the pairwise comparisons between the three developmental stages. Proteins with less than 6 valid values in at least 1 group were filtered out. False positives were excluded utilizing the Benjamini–Hochberg false discovery rate test. STRING (Version 12.0, https://string-db.org/) annotation was used to further characterize DE proteins (Szklarczyk et al. 2023). Venn diagram was generated using the jvenn web application (https://jvenn.toulouse.inrae.fr/app/index.html).

The selection of proteins defined as ’neuronal’ was performed manually by filtering the functional annotations produced by STRING after querying the Gene Ontology databases (Biological Process and Cellular Component), Reactome, InterPro, and STRING clusters for the terms ’neuron*’ and ’nerv*’.

#### Pathway and Gene Ontology enrichment analysis

Functional enrichment analysis was performed with the g:GOSt tool of the g:Profiler web server (https://biit.cs.ut.ee/gprofiler/gost). The enrichment bubble plots were generated using the online SRplot tool (https://bioinformatics.com.cn) and edited with the Adobe Illustrator CS3 software (version 13.0.0).

#### Protein-protein interaction (PPI) network analysis

Network analysis of differentially expressed proteins across the three comparisons was performed using STRING, applying the full network type and a minimum required interaction score of 0.7 (high confidence). Network clustering was performed using the modularity optimization algorithms “multi-level” (Blondel et al. 2008) and “fast greedy” (Clauset, Newman, and Moore 2004), available in the R package igraph (https://doi.org/10.48550/arXiv.2311.10260). The functional cartography was performed using the network analysis tool dmfind v2 (Bersanelli et al. 2016), available at the URL https://github.com/emosca-cnr/dmfind. The participation coefficient (P) and the within-module (community) degree z-score (z) of a protein 𝑖 were defined as:

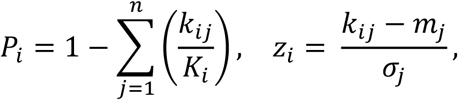

where 𝑛 is the number of communities, 𝐾_𝑖_ is the number of interactions of protein 𝑖, 𝑘_𝑖𝑗_ is the number of interactions of protein 𝑖 within community 𝑗, 𝑚_𝑗_ is the average 𝑘_𝑖𝑗_ over all proteins of the community 𝑗, and 𝜎_𝑗_ is the standard deviation of 𝑘_𝑖𝑗_ over all proteins of the community 𝑗.

The over representation analysis on network communities was performed using protein GO biological process (BP) terms provided by STRING (v12) (Szklarczyk et al. 2023). Only GO BP terms containing at most 500 proteins and at least 3 proteins of the same network community were considered. The reference universe for the hypergeometric test was defined by all the proteins of the network under analysis.

The processed networks files were imported into Cytoscape (version 3.10.3) for graphical visualization of communities and major connector hubs within the network, as well as for mapping the proteins previously annotated as ’neuronal’.

## RESULTS

### Proteomic analysis

In order to identify changes in protein levels during the initial stages of Ciona metamorphosis, when CNS degeneration occurs, protein extracts from larvae (200 pooled individuals) at three different developmental stages, Swimming (SwL, stage 28 of Hotta), Settled (SetL, stage 32 of Hotta) and Metamorphosis (MetL, stage 34 of Hotta), were analyzed by liquid chromatography mass spectrometry (LC-ESI-MS/MS). For each stage, three biological replicates were analyzed. After performing quality control on the obtained results, a sample from the SetL stage was discarded due to low or absent signal, likely resulting from technical issues during protein extraction.

Overall, the analysis led to the identification of 1152 proteins. Among them, 405 are differentially expressed (DEPs), either up or downregulated, in at least one stage comparison (ANOVA and Tukey’s test, p-value < 0.05) (Table S1). The number of significantly upregulated proteins was 70, 124 and 74 for the SetL versus SwL, MetL versus SetL and MetL versus SwL comparisons, respectively; while 152, 100 and 154 proteins resulted downregulated in the same comparisons (Table 1). A total of 35 proteins, out of 405, were found to be differentially modulated across all the stage comparisons (Figure 1).

**Figure 1.**
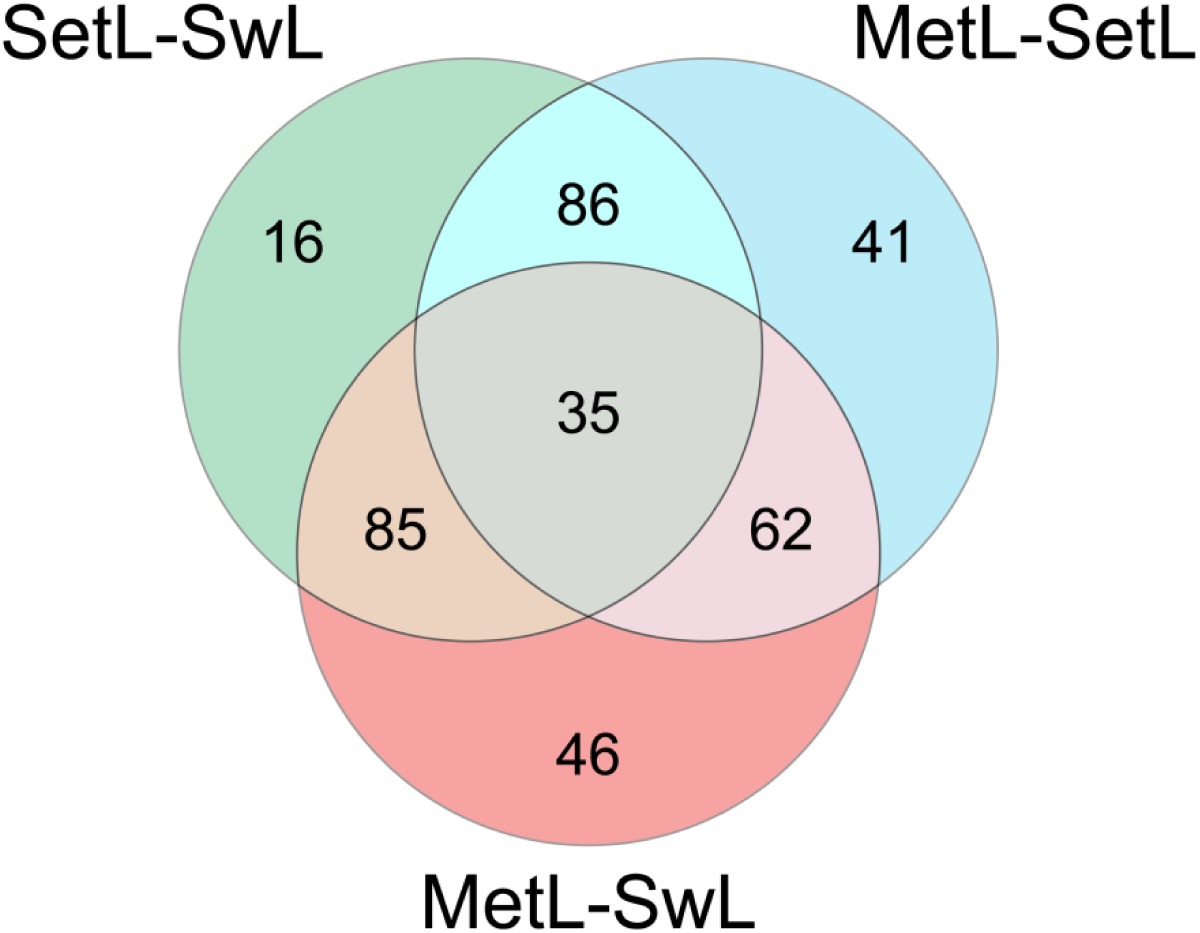
Venn diagram showing the distribution of differentially abundant proteins across the three experimental comparisons. The overlaps represent shared proteins among the comparisons SetL-SwtL, MetL-setL, and MetL-SwL, while unique regions indicate proteins specifically varying in a single comparison. Numbers within each section correspond to the count of proteins in that category.

**Table 1.** Number of overexpressed and underexpressed proteins in the three stages comparisons.

|  | **Overexpressed** | **Underexpressed** | **Total** |
| --- | --- | --- | --- |
| **SetL vs. SwL** | 70 | 152 | 222 |
| **MetL vs. SetL** | 124 | 100 | 224 |
| **MetL vs. SwL** | 74 | 154 | 228 |

### Annotation of DE proteins

DEPs were then identified integrating the current UniProtKb annotation of Ciona proteome (UP000008144) and the STRING annotation (Table S1). 283 proteins were fully annotated with the corresponding protein name and associated molecular function/activity, while other 99 were classified based upon the presence of a specific domain (referred to as ‘domain-containing protein’). The most represented domains are VWFA domain, EF-hand domain, RRM domain, IF rod domain and Sushi domain. Finally, 23 proteins remain completely uncharacterized (‘uncharacterized protein’). The annotation of the top 10 overexpressed and top 10 underexpressed proteins in the three stage comparisons is shown in Tables 2, 3, and 4, respectively. Notably, several annotated DEPs displayed an opposite regulation when comparing the transition between the SwL and SetL stage towards the transition between the SetL and MetL stage, with 32 of them being upregulated going from the first to the second stage and downregulated from the second to the third stage, while 70 showing an inverse trend (see Table 5 and S1). We also incorporated an additional layer of annotation by cross-referencing DEPs with annotations indicating relevance to the nervous system and neuronal functions, as supported by multiple data sources (Table S2). We identified 21, 23 and 32 neuronal DEPs for the SetL-SwL, MetL-SetL and MetL-SwL comparisons, respectively.

**Table 2.** List of the top 10 overexpressed and 10 underexpressed proteins in the SetL-SwL comparison.

| Rank | UniprotKB ID | Protein name<sup>a</sup> | Gene name<sup>b</sup> | Log2FC SetL/SwL |
| --- | --- | --- | --- | --- |
| **Overexpressed** |  |  |  |  |
| 1 | F6Q7V3 | VWFA domain-containing protein | na | 3.718 |
| 2 | F6YF25 | VWFA domain-containing protein | na | 1.949 |
| 3 | F6R9M9 | EF-hand domain-containing protein | na | 1.938 |
| 4 | H2Y0U3 | Uncharacterized protein | na | 1.934 |
| 5 | F6V1M6 | Uncharacterized protein | meta2 | 1.868 |
| 6 | H2XLP7 | VWFA domain-containing protein | na | 1.757 |
| 7 | F6QFI4 | Signal peptidase complex subunit 2 | LOC100178515 | 1.626 |
| 8 | F6R1X5 | Cell division cycle 42 | cdc42 | 1.619 |
| 9 | F6XQP8 | ZnF_CDGSH domain-containing protein | LOC100179954 | 1.434 |
| 10 | F6R9G4 | EF-hand domain-containing protein | na | 1.424 |
| **Underexpressed** |  |  |  |  |
| 1 | F6QHG3 | Uncharacterized protein | na | -3.006 |
| 2 | F6YX55 | VWFA domain-containing protein | na | -2.272 |
| 3 | F6U9F4 | Cytoplasmic aconitate hydratase | LOC100186938 | -2.174 |
| 4 | H2Y1H9 | Protein transport protein SEC23 | LOC100183335 | -1.917 |
| 5 | F7AJ84 | Glucosidase II subunit alpha | na | -1.770 |
| 6 | F6WUH0 | NADH dehydrogenase [ubiquinone] iron-sulfur protein 3, mitochondrial (Complex I-30kD) | LOC100183052 | -1.696 |
| 7 | F7AKV1 | Villin-1 | LOC100185031 | -1.681 |
| 8 | F6X0F3 | Microtubule-associated protein RP/EB family member 1 | LOC100181072 | -1.666 |
| 9 | F6PTZ3 | Endothelin-converting enzyme 1 | na | -1.598 |
| 10 | F6U8C7 | Eukaryotic translation initiation factor 3 subunit B (eIF3b) | LOC100182767 | -1.595 |

<sup>a</sup> UniProtKB annotation - release 2024\_04<sup>b</sup> NCBI Gene

**Table 3.**
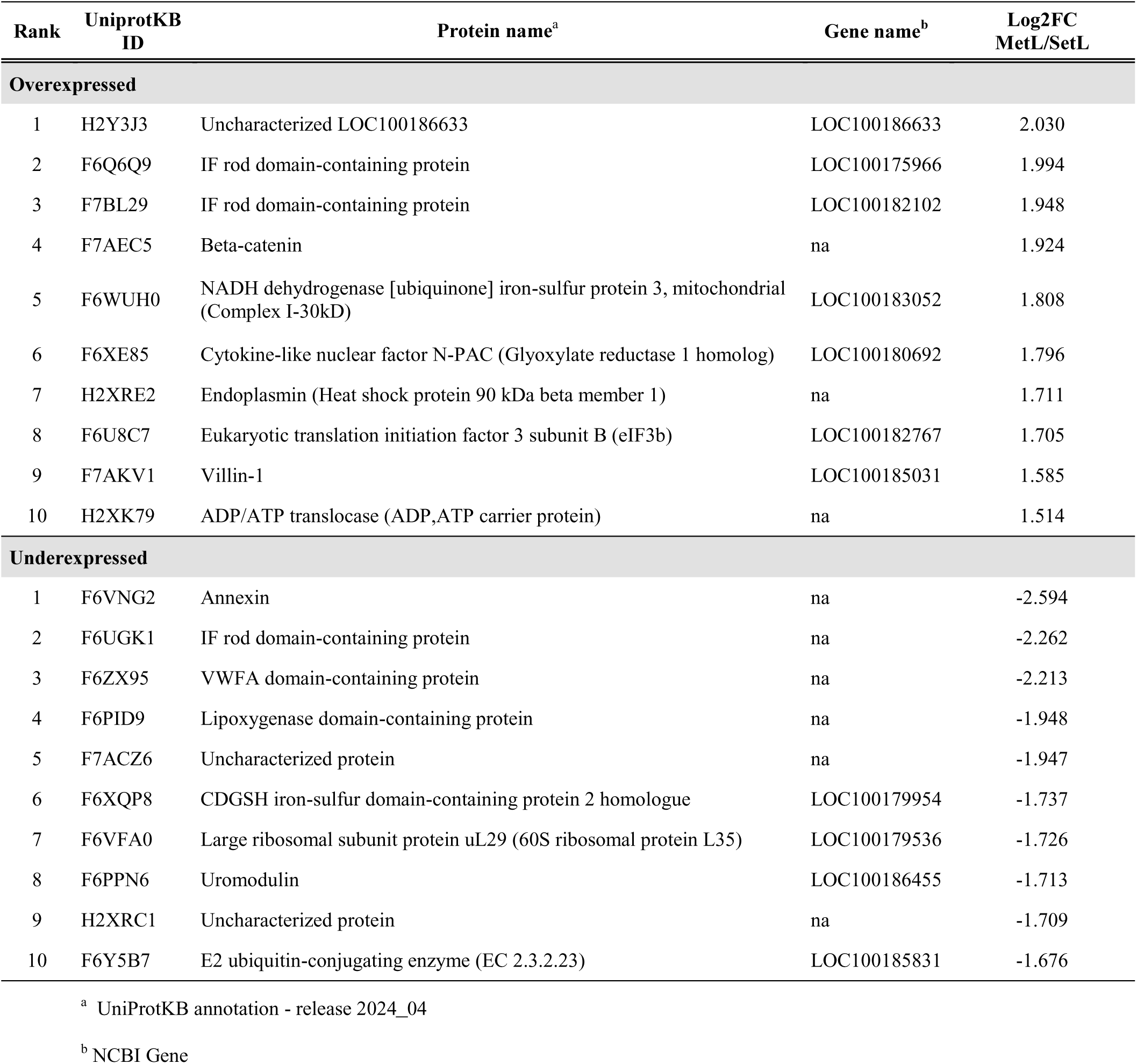
List of the top 10 overexpressed and 10 underexpressed proteins in the MetL-SetL comparison.

**Table 4.**
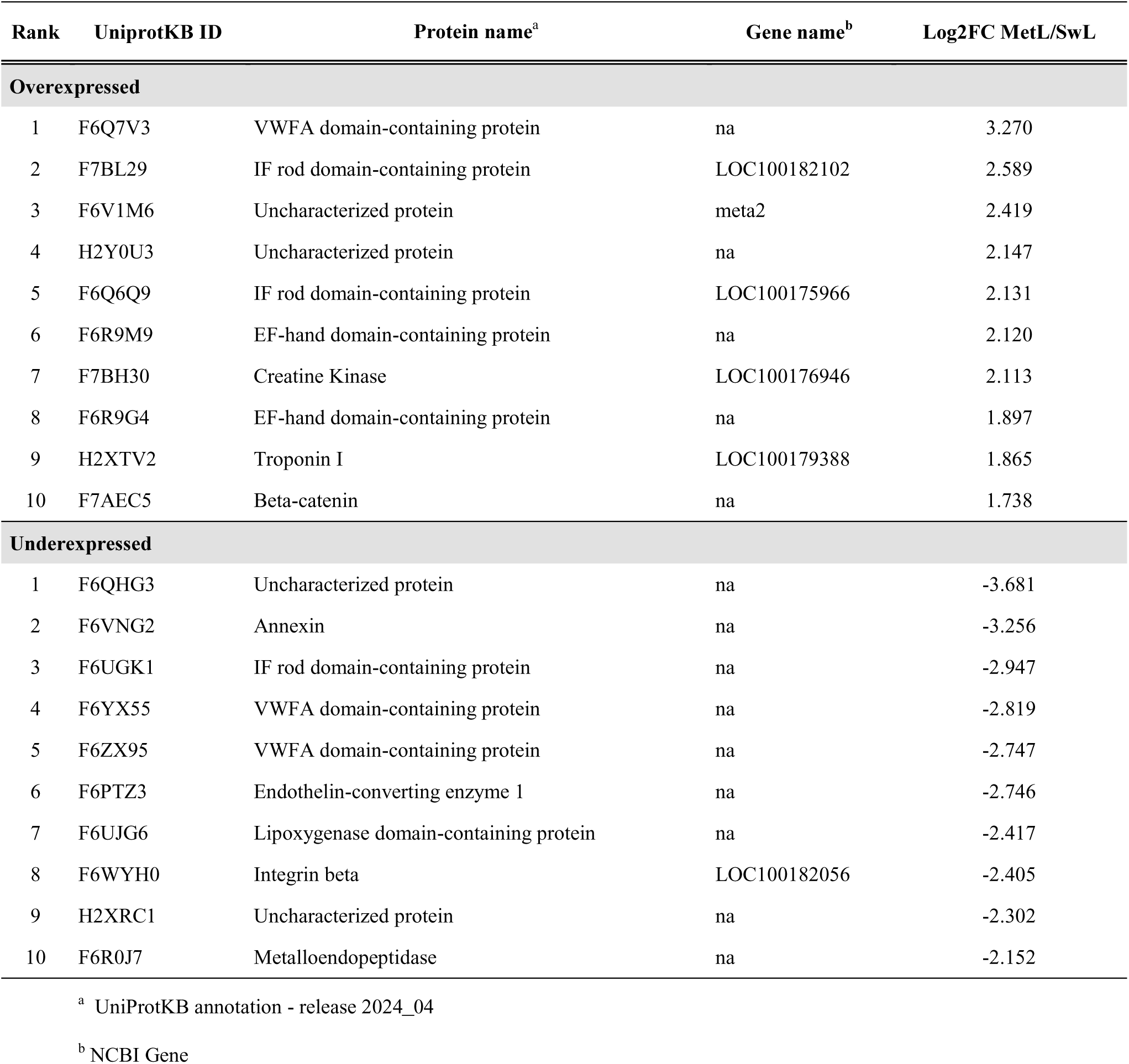
List of the top 10 overexpressed and 10 underexpressed proteins in the MetL-SwL comparison.

**Table 5.**
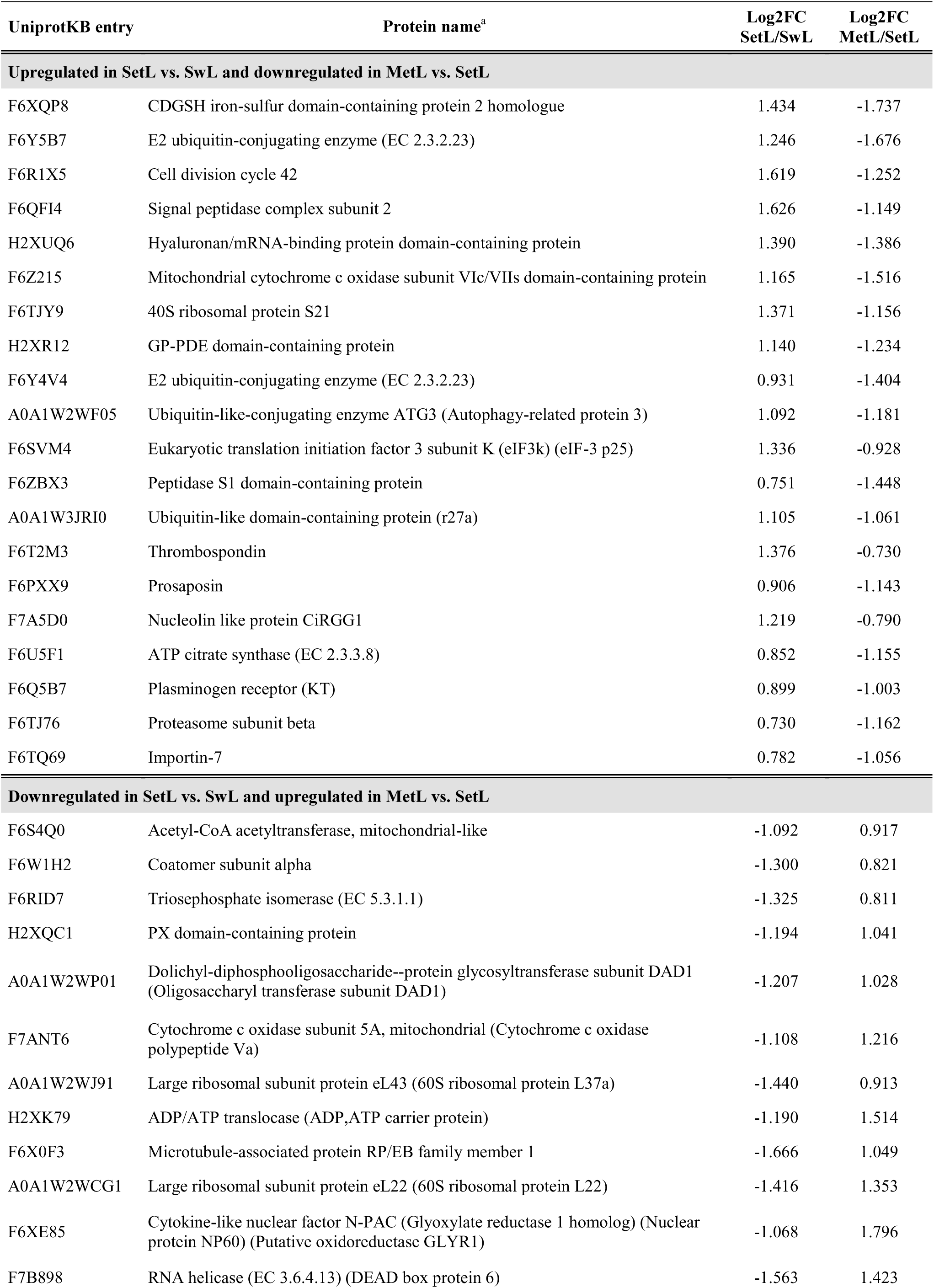
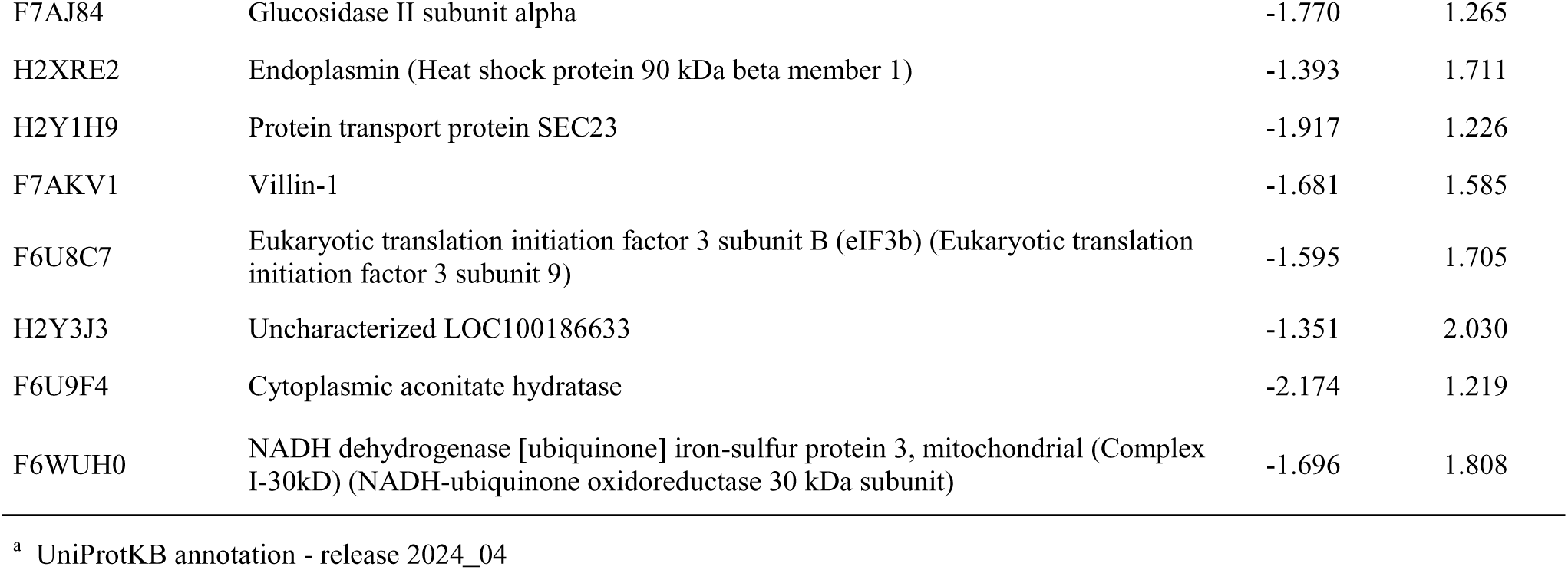
List of the most significant proteins displaying opposite regulation in the SetL-SwL and MetL-SetL comparisons.

### Enrichment analysis

In order to gain insights into the pathways and biological processes that are dynamically regulated during the metamorphosis, we carried out a KEGG pathway and Gene Ontology enrichment analysis on DEPs (Figure 2, S1-2; Tables S3-5). The analysis revealed enriched KEGG terms that were common to all stage comparisons, including Motor proteins, Oxidative phosphorylation, Fatty acid degradation and Biosynthesis of amino acids. On the contrary, some pathways were found to be specifically enriched only in one comparison. Notably, Autophagy pathway (comprising an autophagy-related gene and mitogen-activated protein kinases), mTOR signaling pathway, Proteasome pathway (including proteasome and proteasome regulatory subunits) and Glyoxylate and dicarboxylate metabolism were significantly enriched during the settlement of the swimming larva (SetL vs SwL). An enrichment in Citrate cycle (TCA cycle), Valine, leucine and isoleucine degradation, Propanoate metabolism and 2-Oxocarboxylic acid metabolism was found during the transition between the settled (SetL) and the metamorphosing (MetL) larva. Finally, pathways related to the metabolism and biosynthesis of some amino acids (particularly arginine, phenylalanine, tyrosine and tryptophan) and Endocytosis (including proteins involved in the actin cytoskeleton organization) were enriched going from the SwL to the MetL stage (Figure 2).

**Figure 2.**
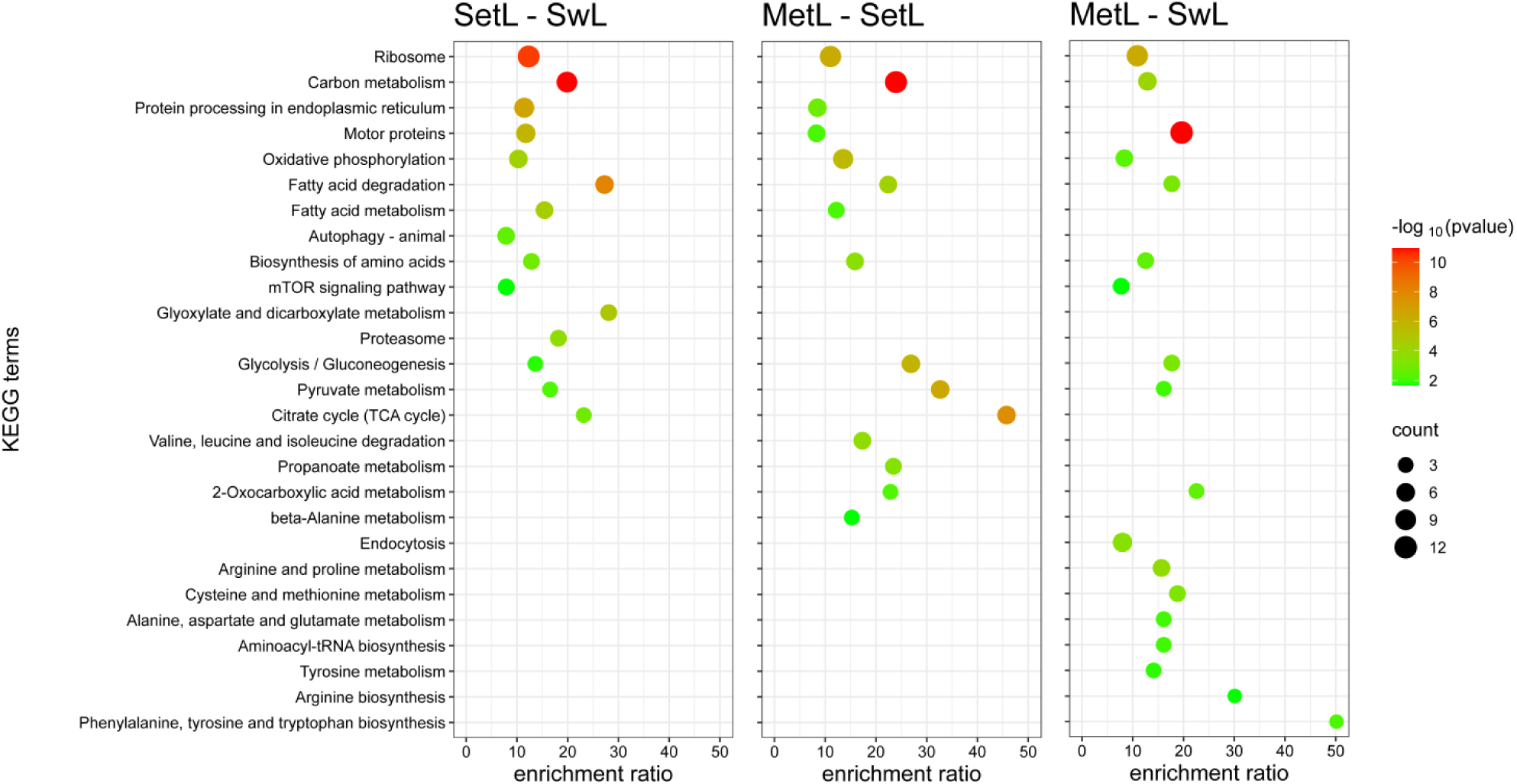
KEGG terms enrichment analysis of differentially expressed proteins across the experimental comparisons. The dot plots show enriched KEGG terms for the comparisons SetL vs SwL, MetL vs SetL, and MetL vs SwL. The x-axis indicates the enrichment ratio, while the y-axis lists the enriched KEGG terms. The color gradient represents the statistical significance (−log₁₀ p-value), with red indicating higher significance, and the dot size corresponds to the number of proteins associated with each term.

Gene Ontology analysis also revealed an involvement of fatty acid and lipid modification, as well as RNA/mRNA binding, for the swimming to settled larva transition. The switch between the SetL and MetL stage, on the other hand, was enriched in GO terms associated to nucleotide and carbohydrate metabolism/catabolism, tricarboxylic acid cycle, and regulation of translation. An enrichment in biological processes and molecular functions related to cytoskeleton organization (particularly actin cytoskeleton) was evidenced for the MetL vs SwL comparison (Figure S1-2).

### Network analysis

To further uncover relevant functional modules and hub proteins involved in the metamorphosis of Ciona during neurodegeneration, we carried out a protein-protein interaction (PPI) network analysis. For each comparison, we defined a network among DEPs using the PPI from the STRING database (https://string-db.org/) (Szklarczyk et al. 2023). Then, we identified network communities (clusters of proteins that exhibit a tendency to interact with one another) and obtained the so-called functional cartography (Guimerà and Amaral 2005), a classification of proteins into roles according to their pattern of intra- and inter-community interactions. This method allowed us to distinguish between hub proteins and non-hubs proteins of different kinds, and their relationships in each comparison (Figure 3, S3-5; Table 6, S6-11).

**Figure 3.**
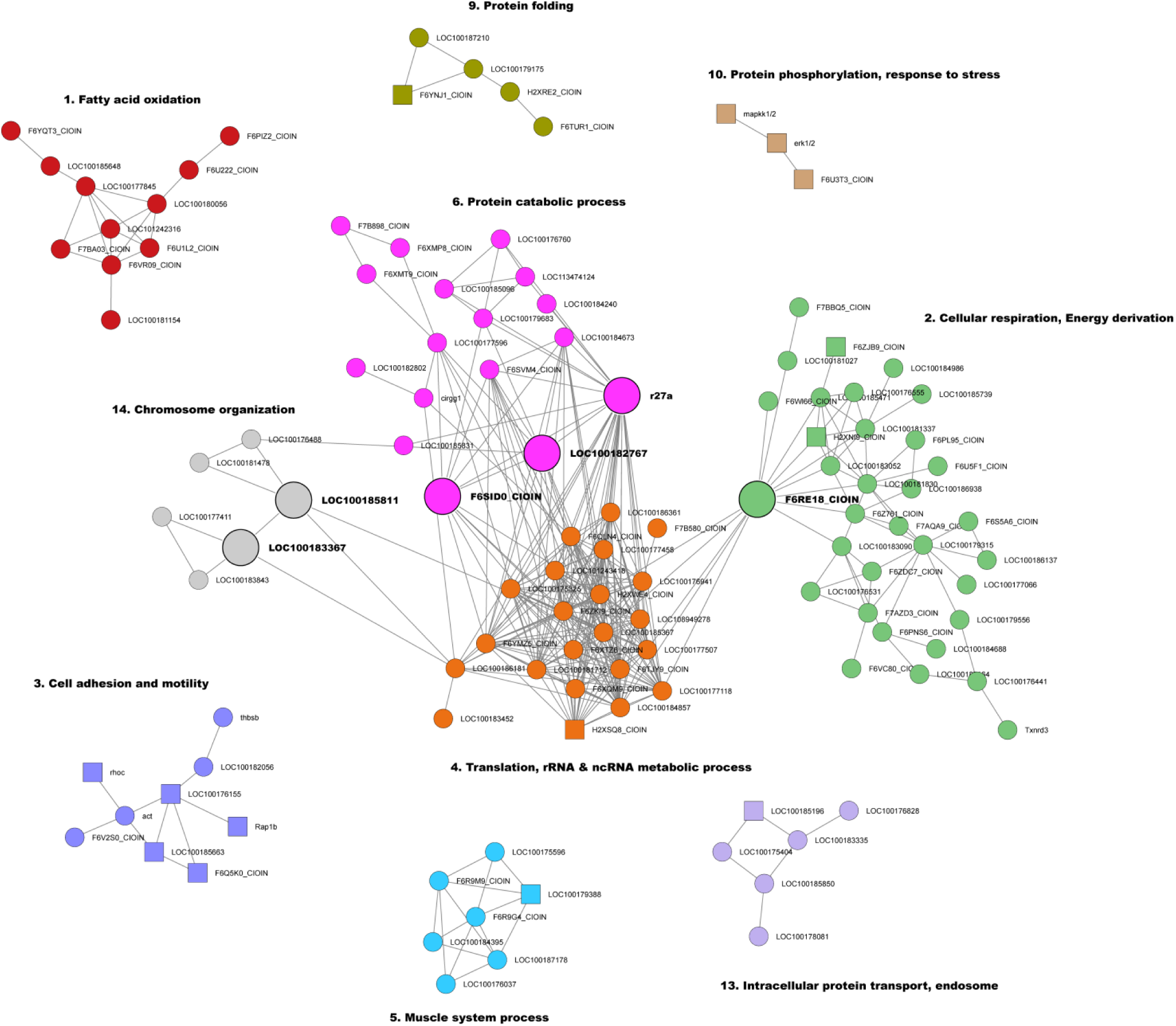
Network analysis of differentially expressed proteins in the SetL-SwL comparison. Node colors correspond to functional clusters (communities), and lines represent connections between proteins. Node sizes indicate the main connector hubs (see Table 6). Square nodes represent proteins annotated as “Neuronal” (see Suppl. Table S2). Major clusters are annotated with the most representative biological process names, highlighting key functional modules.

**Table 6.**
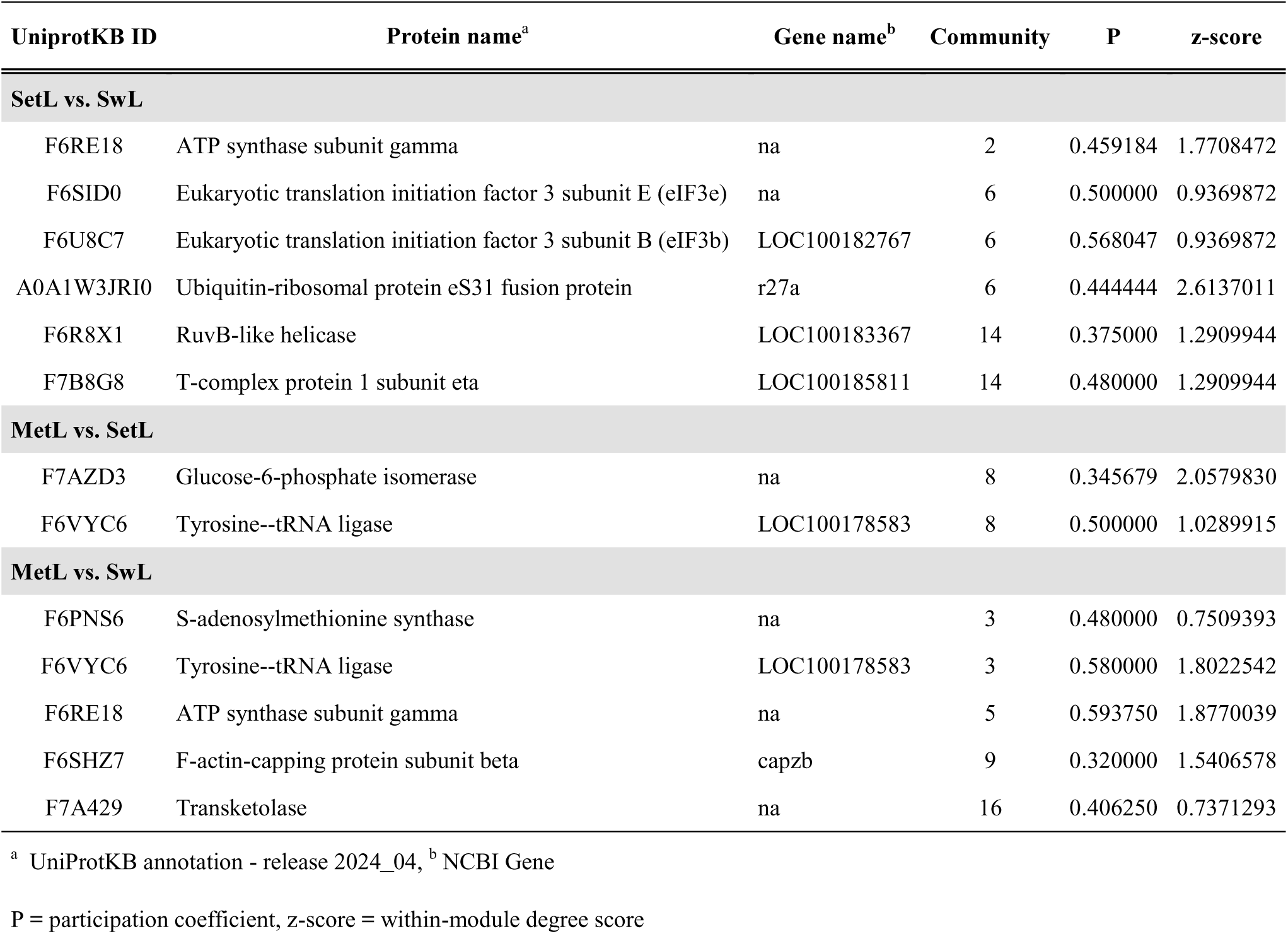
List of connector hub proteins identified by PPI Network Analysis.

For the transition between the SwL and SetL stage, a total of 15 modules (communities) were identified, 10 of which contained at least three proteins (Figure 3). Notably, four of these clusters are interconnected through six hub proteins (‘connector hubs’) (Table 6, Figure S5). Modules 4, enriched for translation regulation and rRNA metabolic process, and 6, related to protein catabolic process, are tightly connected through two subunits of Eukaryotic translation initiation factor (eIF3b and eIF3e) and the Ubiquitin-ribosomal protein eS31 fusion protein (r27a). These two modules are also linked to module 14 (chromosome organization) thanks to RuvB-like helicase and T-complex protein 1 subunit eta. Proteins belonging to Module 2, involved in cellular respiration, energy derivation and metabolism of nucleotides, and 4 (translation) are connected by ATP synthase subunit gamma. Modules 3 (cell adhesion and motility) and 10 (protein phosphorylation and response to stress) are particularly enriched in proteins involved in CNS functions (Figure 3). The MetL-SetL network is characterized by 19 clusters, 12 with at least three proteins. Seven of these modules are interconnected, with two hub proteins, Glucose-6-phosphate isomerase and Tyrosine-tRNA ligase, linking carbohydrate metabolism (Module 8) to fatty acid oxidation and cellular respiration processes (modules 2 and 19), on the one hand, and to translation, rRNA processing, protein folding and stabilization (Modules 3, 5 and 12), on the other hand (Figure S3 and S5, Table 6). Module 1 is enriched with proteins involved in organismal development and neurogenesis. Finally, by analyzing the whole metamorphosis process, from SwL to MetL stage, we identified 19 modules (16 containing at least three proteins). 8 modules are interconnected through the interaction of five connector hub proteins (Figure S4-5, Table 6). Central to this network is the Transketolase enzyme, bridging together carbohydrate metabolism (Module 16), tRNA metabolic process (Module 3, hub proteins S-adenosylmethionine synthase and Tyrosine--tRNA ligase), cellular respiration (Module 5, hub protein ATP synthase subunit gamma) and actin cytoskeleton organization (Module 9, hub protein F-actin-capping protein subunit beta). Of note, Modules 1, related to cell adhesion and neurogenesis, and 9 (actin cytoskeleton organization) contain several proteins annotated as ‘neuronal’.

## DISCUSSION

We employed a proteomic strategy to identify novel factors potentially involved in neurodegeneration, leveraging the ascidian Ciona, which naturally undergoes a programmed degeneration of its nervous system as the swimming larva transitions into a sessile juvenile. This approach is well supported by prior studies, as proteomic analyses have proven powerful in revealing molecular changes underlying neurodegenerative disorders in humans (Moya-Alvarado et al. 2016). Moreover, in Ciona proteomic analysis already proved to be a successful approach to quantify proteins in unfertilized eggs and to track proteomic changes throughout embryogenesis (Frese et al. 2024).

We identified 1,152 proteins, among them 405 are differentially expressed in at least one stage comparison and could be important tools to delineate the changes at cellular level occurring during the metamorphosis of Ciona. Most of the differentially expressed proteins (DEPs) were identified and associated with their function, using the most updated proteome annotation of Ciona.

Interestingly, several DEPs are annotated as containing domains characteristic of proteins involved in cellular processes that play key roles during development, such as cell adhesion (VWFA), calcium signaling (EF-Hand) and cytoskeleton assemblage (IF rod). Other DEPs were identified as Sushi domain-containing proteins. Despite the function of many proteins containing this domain remains unknown, their role in neural synapsis regulation has been suggested in different animals (Gendrel et al. 2009; M. Nakayama et al. 2016) and thus their involvement in the remodeling of nervous system in Ciona during metamorphosis can be hypothesized.

Remarkably, several DEPs identified in our analysis have also been reported as dysregulated or impaired in patients with Alzheimer’s disease (Moya-Alvarado et al. 2016) or other neurodegenerative diseases, including Annexin, Cdc42, Glial fibrillary acidic protein, NADH dehydrogenase, Peptidyl-prolyl cis-trans isomerase and Transketolase, among others (some relevant examples are reported in Table 7) (Bartolome et al. 2020; Bedrood et al. 2009; Ries et al. 2016; Davis et al. 2009; Saraceno et al. 2018; Umbayev et al. 2023; Zhu et al. 2023; Al Mamun et al. 2020; Chen et al. 2025; Kawahata and Fukunaga 2023; Leipp et al. 2024; Keeney et al. 2006; Marella et al. 2009; Lee, Pastorino, and Lu 2011; Xu et al. 2017; He et al. 2023; Oji et al. 2020; Gibson et al. 2000; Vinh Quc Lu’O’Ng and Nguyen 2011). This observation suggests that common mechanisms may underlie pathological neurodegeneration in humans and physiological neurodegeneration in ascidians and further supports the validity of our approach.

**Table 7.** Examples of Ciona DEPs whose human orthologue is involved in neurodegenerative diseases.

| Protein name | Role in neurodegenerative diseases | References |
| --- | --- | --- |
| Annexin | Anti-inflammatory role; role in the prevention and reduction of amyloid-induced toxicity. | Bartolome et al. 2020; Bedrood et al. 2009; Ries et al. 2016 |
| Capz | Decreased levels are associated with neurites and growth cones defects. | Davis et al. 2009 |
| Cell division cycle 42 | Upregulated in AD with associated loss of dendritic spines; upregulated in FTLD. | Saraceno et al. 2018; Umbayev et al. 2023; Zhu et al. 2023 |
| E2 ubiquitin-conjugating enzyme | Impaired function allows Aβ and tau to accumulate, forming plaques and tangles in AD; involved in early synaptic dysfunction. | Al Mamun et al. 2020 |
| Fatty acid-binding protein | Involved in α-synuclein aggregation, neurotoxicity, and neuroinflammation in PD; involved in neuroinflammation in AD. | Chen et al. 2025; Kawahata and Fukunaga 2023 |
| Glial fibrillary acidic protein | Upregulated and involved in astrogliosis in AD and other neurodegenerative diseases. | Leipp et al. 2024 |
| NADH dehydrogenase | Decreased activity linked to mitochondrial impairment, oxidative stress and neuron loss in PD. | Keeney et al. 2006; Marella et al. 2009 |
| Peptidyl-prolyl cis-trans isomerase | Reduced activity linked to Aβ and tau pathology and loss of synaptic plasticity in AD. | Lee et al. 2011; Xu et al. 2017 |
| Prosaposin | Involved in lipid homeostasis, protective role in PD; mutations associated to PD. | He et al. 2023; Oji et al. 2020 |
| Transketolase | Impaired TK activity reduces neurogenesis, contributing to cognitive dysfunction in AD and other neurodegenerative conditions. | Gibson et al. 2000; Vinh Quc Lu'O'Ng and Nguyen 2011 |

AD = Alzheimer's Disease; FTLD = Frontotemporal Lobar Degeneration; PD = Parkinson's Disease

Another validation of our results was the identification of the Meta-2 protein among the top ten upregulated ones in the comparison between swimming larvae and settled larvae. This observation is consistent with a transcriptome analysis in Ciona, which reported that *Ci-meta2* expression begins at the larval stage and is upregulated in metamorphosing juveniles (A. Nakayama, Satou, and Satoh 2001). *Ci-meta2* expression was detected in the adhesive organs, structures involved in settlement, as well as in the neck region of the central nervous system and in the dorsal epidermis of the trunk corresponding to the siphon primordium. It has been suggested that Meta-2 may play a role in the dynamic rearrangement of cells during ascidian metamorphosis (A. Nakayama, Satou, and Satoh 2001).

Our analysis revealed that numerous DEPs are specifically upregulated or downregulated in the settled larva when compared with both the swimming larva and the metamorphosed juveniles. These patterns point to a unique proteomic landscape characterizing the settled stage, in which substantial molecular remodeling occurs and critical pathways are differentially regulated relative to adjacent developmental stages. This is especially relevant to our study, as neurodegeneration is initiated and reaches its peak during this stage (Hotta, Dauga, and Manni 2020; Comes et al. 2007). Proteins with this peculiar expression profile could be candidate players of the physiological neurodegeneration process.

We observed that several proteins with this expression profile are involved in autophagy, ubiquitination, and proteasome regulation. Among these, r27a resulted also a hub protein in network analysis linking functional modules of catabolic process and those of translation, rRNA and scRNA metabolic process. It is well known that autophagy is a crucial process for embryonic and postnatal development, contributing to cell fate decisions by removing components that are no longer needed, which is critical for proper differentiation. mTOR (mammalian target of rapamycin) is a key actor of autophagic process and Gene Ontology enrichment analysis evidenced that its signaling pathway was significantly upregulated in SetL compared to both SwL and MetL stages. mTOR is a protein kinase that regulates cell growth, metabolism, proliferation, and survival. It is a master regulator that integrates signals from nutrients, growth factors, and energy levels to control a cell’s growth and functions. Dysregulation of the mTOR pathway is implicated in diseases like cancer, diabetes, and aging (Schmeisser and Parker 2019; Perluigi, Di Domenico, and Butterfield 2015). The ubiquitin-proteasome system (UPS) targets cellular proteins for degradation. Defects in the UPS have been found to play a crucial role in the pathogenesis of Alzheimer’s disease (AD). Current investigations have revealed that UPS components are associated with both the early stage of AD, which is characterized by synaptic dysfunction, and the late stages of the disease, which are marked by neurodegeneration (Al Mamun et al. 2020).

Enrichment and network analyses both evidenced that the overall analyzed process, from swimming larva to settled larva, is characterized by differential expression of proteins involved in actin cytoskeleton organization. Noteworthy, the specific community (actin cytoskeleton organization) in network analysis contains several proteins annotated as ‘neuronal’ such as RapB1, a GTP-binding protein, and RhoC, a small GTPase that plays a key role in cell migration, cytoskeletal organization, and cell division. Notably, abnormalities in the neuronal cytoskeleton are characteristic of several neurodegenerative disorders (Wurz et al. 2022). Among DEPs detected by our analysis correlated with actin cytoskeleton, Cdc42, Capz and Villin-1 are of particular interest.

Cdc42 (Cell division cycle 42) protein is among the top 10 overexpressed proteins in the transition from SwL to SetL stage, and it is subsequently downregulated from SetL to MetL (Table 2 and 5). Cdc42 is a member of the Rho GTPase family that participates in axonogenesis and is known to play a role in the development and progression of neurodegenerative disorders (Umbayev et al. 2023). Members of this family are regulators of F-actin polymerization (Bishop and Hall 2000), acting as molecular switches by cycling between an inactive GDP-bound state and an active GTP-bound state.

Dysregulation of Cdc42 seems to be a shared trait of different neurodegenerative conditions. The progressive loss in the number of dendritic spines characteristic of Alzheimer’s disease (AD) has been correlated with Cdc42 activity upregulation (Zhu et al. 2023). Increased levels of Cdc42 were also reported in the frontal cortex of patients with frontotemporal lobar degeneration (FTLD), compared to age-matched controls (Saraceno et al. 2018). Moreover, Cdc42-interacting protein 4 (CIP4) is a Cdc42 effector protein that is involved in cytoskeletal organization and accumulates with neuropathological severity in the neostriatum of Huntington’s disease (HD) patients. It associates with Huntingtin (HTT) and its accumulation and cellular toxicity may have a role in HD pathogenesis (Holbert et al. 2003).

Noteworthy, both alpha and beta subunits of Capz, another actin cytoskeleton interacting protein, are downregulated during Ciona metamorphosis and were found to be hub proteins in the MetL-SwL network (Figure S4, Table 6 and S1). Capz is an actin capping protein that regulates actin filament growth by binding to the barbed ends of actin filaments, thereby preventing further polymerization. In neurons, capz is critical for normal neurite outgrowth and growth cone morphology. Its silencing causes short, dystrophic neurites reminiscent of the cytoskeletal defects seen in neurodegenerative diseases. Capz role in regulating actin dynamics and microtubule polymerization through interactions with tau protein links it to neuronal morphology and network stability, which are disrupted during neurodegeneration (Davis et al. 2009; Fulga et al. 2007). Cdc42 and Capz in neurodegeneration have opposing but complementary roles in actin cytoskeleton remodeling. While cdc42 activation promotes actin polymerization and filopodia formation, processes essential for dendritic spine maintenance and neuronal connectivity, Capz caps actin filaments to regulate and stabilize actin networks. Disruption of this balance can lead to aberrant actin dynamics, neurite dystrophy, and synaptic loss, contributing to the progression of neurodegenerative disorders (Zhu et al. 2023; Davis et al. 2009).

Another protein interacting with actin cytoskeleton that is upregulated in SetL stage compared to previous and following stages is Villin-1. It is a major actin-modifying protein that is associated with the microvillar actin filaments. In homo, it is expressed prevalently in renal and gastrointestinal epithelial cells. It has been demonstrated that Villin regulates epithelial cell morphology, actin reorganization, and cell motility (Tomar et al. 2006). Villin has a pro-apoptotic role and its high levels in the swimming larva can be preparatory for the apoptosis that characterized the transition toward the settled larva. Interestingly, another member of the Villin-1 family, Villin-2 (Ezrin), a member of the ezrin/radixin/moesin (ERM) family of actin interacting proteins, was identified as a protein that is upregulated in tau-mediated neurodegeneration in mice and in the temporal cortex of mild cognitive impairment (MCI) and AD subjects. Ezrin protein abundance changes precede motor impairment and are associated with the early stages of neurodegeneration in human disease (Vega et al. 2018).

Several other proteins involved in Citrate cycle (TCA cycle), Valine, leucine and isoleucine degradation, Propanoate metabolism and 2-Oxocarboxylic acid metabolism, were found overexpressed during the transition between the settled and the metamorphosing larva, indicating activation of metabolic processes necessary to rebuild the juvenile organs.

A hub protein bringing together different identified modules in the comparison between SwL and MetL stages is the enzyme transketolase (TK) (Figure S4, Table 6). TK deregulation is strongly connected to neurodegeneration and neurodegenerative diseases in humans primarily through its role in thiamine (vitamin B1) metabolism and the pentose phosphate pathway (PPP). TK is a thiamine-dependent enzyme critical for the PPP, where it facilitates the synthesis of NADPH and ribose-5-phosphate, essential for maintaining cellular redox balance and nucleotide biosynthesis, respectively (Zhao et al. 2014). In conditions of thiamine deficiency, TK activity is markedly reduced, which impairs the PPP. This leads to decreased production of NADPH and ribose-5-phosphate, crucial for combating oxidative stress and for neurogenesis, particularly in the hippocampus. Impaired TK activity reduces hippocampal neurogenesis, inhibits proliferation of hippocampal progenitor cells, and affects differentiation into neurons, including cholinergic neurons. This contributes to cognitive dysfunction observed in neurodegenerative conditions such as Wernicke’s encephalopathy and Alzheimer’s disease (Vinh Quc Lu’O’Ng and Nguyen 2011; Gibson et al. 2000).

Our network analysis also identified different communities associating proteins with recognized function and unknown proteins. Some of these communities included proteins that have been recognized as “neural” (Figure 3 and S3-4, Table S2). We think that further analysis of components of these communities could be promising to identify new actors in neurodegeneration.

We are aware that our study have some limitations. First, we identified 1,152 proteins, representing only a subset of the 7,057 unique proteins reported by Frese et al. (Frese et al. 2024) that tracked proteomic changes throughout embryogenesis of Ciona, when at least half of the protein-coding genes are expressed. Although our analysis encompassed only three developmental stages, we cannot rule out that the relatively low number of detected proteins can be due to technical limitations. Moreover, we analyzed the proteome of the entire organisms and we could have missed more subtle changes specifically occurring in the nervous system. Furthermore, the incomplete functional annotation of proteins in this species hampers a comprehensive interpretation of the underlying biological processes. Nevertheless, the set of DEPs and processes identified here provides a valuable basis for elucidating the main mechanisms involved in ascidian neurodegeneration. Importantly, this study revealed an unexpected similarity with the neurodegeneration process in human, and suggested that some molecular mechanisms at the basis of neurodegeneration are remarkably conserved throughout the chordate evolution. This observation further supports ascidians as valuable models to study neurodegenerative diseases.

## AUTHOR CONTRIBUTION

DC, EM: Data curation, Formal analysis, Visualization. MV, DC, SM, RP: Writing-review & editing, Conceptualization, Investigation. SM, RP, DC, CB, EM: Writing-review & editing, Methodology. RP, MV, DC: Conceptualization, Supervision, Funding acquisition.

## ACKNOWLEDGMENTS

The authors acknowledge the Unitech OMICS platform at the University of Milan for Orbitrap LC-MS/MS analysis. The authors acknowledge the student Alberto De Benedectis for supporting the first phase of the proteome data analysis during its Master degree internship.

## DATA AVAILABILITY

The data supporting the findings of this study are available within the article and its supplemental data. Raw data that support the findings of this study are openly available in the UNIMI Dataverse repository with persistent identifier number: doi.org/10.13130/RD_UNIMI/K091MC, reference name “SEED_IMPATTO_PROGETTO”.

## SUPPORTING INFORMATION

**Table S1.** Annotated list of differentially expressed proteins in at least one stage comparison.

**Table S2.** List of differentially expressed proteins annotated as involved in neuronal functions for the three stage comparisons.

**Table S3.** Gene Ontology and KEGG pathway enrichment analysis carried out on the differentially expressed proteins identified in the SetL-SwL comparison.

**Table S4.** Gene Ontology and KEGG pathway enrichment analysis carried out on the differentially expressed proteins identified in the MetL-SetL comparison.

**Table S5.** Gene Ontology and KEGG pathway enrichment analysis carried out on the differentially expressed proteins identified in the MetL-SwL comparison.

**Table S6.** Functional cartography analysis performed on SetL-SwL network.

**Table S7.** Functional cartography analysis performed on MetL-SetL network.

**Table S8.** Functional cartography analysis performed on MetL-SwL network.

**Table S9.** GO biological process (BP) over representation analysis on protein communities identified in the SetL-SwL network.

**Table S10.** GO biological process (BP) over representation analysis on protein communities identified in the MetL-SetL network.

**Table S11.** GO biological process (BP) over representation analysis on protein communities identified in the MetL-SwL network.

**Figure S1.**
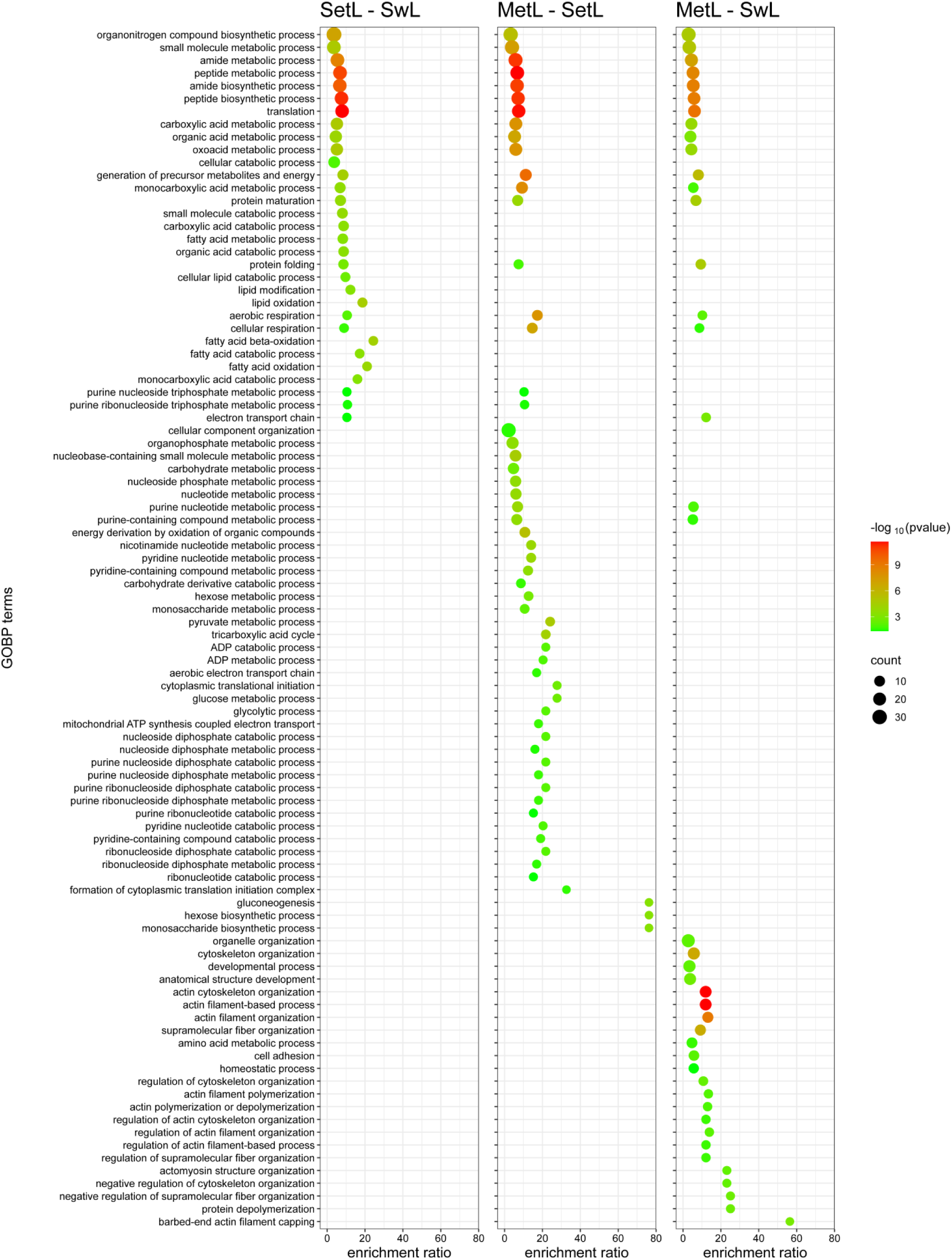
Gene Ontology Biological Process (GOBP) terms enrichment analysis of differentially expressed proteins across the experimental comparisons. The dot plots show enriched GOBP terms for the comparisons SetL vs SwL, MetL vs SetL, and MetL vs SwL. The x-axis indicates the enrichment ratio, while the y-axis lists the enriched GOBP terms. The color gradient represents the statistical significance (−log₁₀ p-value), with red indicating higher significance, and the dot size corresponds to the number of proteins associated with each term.

**Figure S2.**
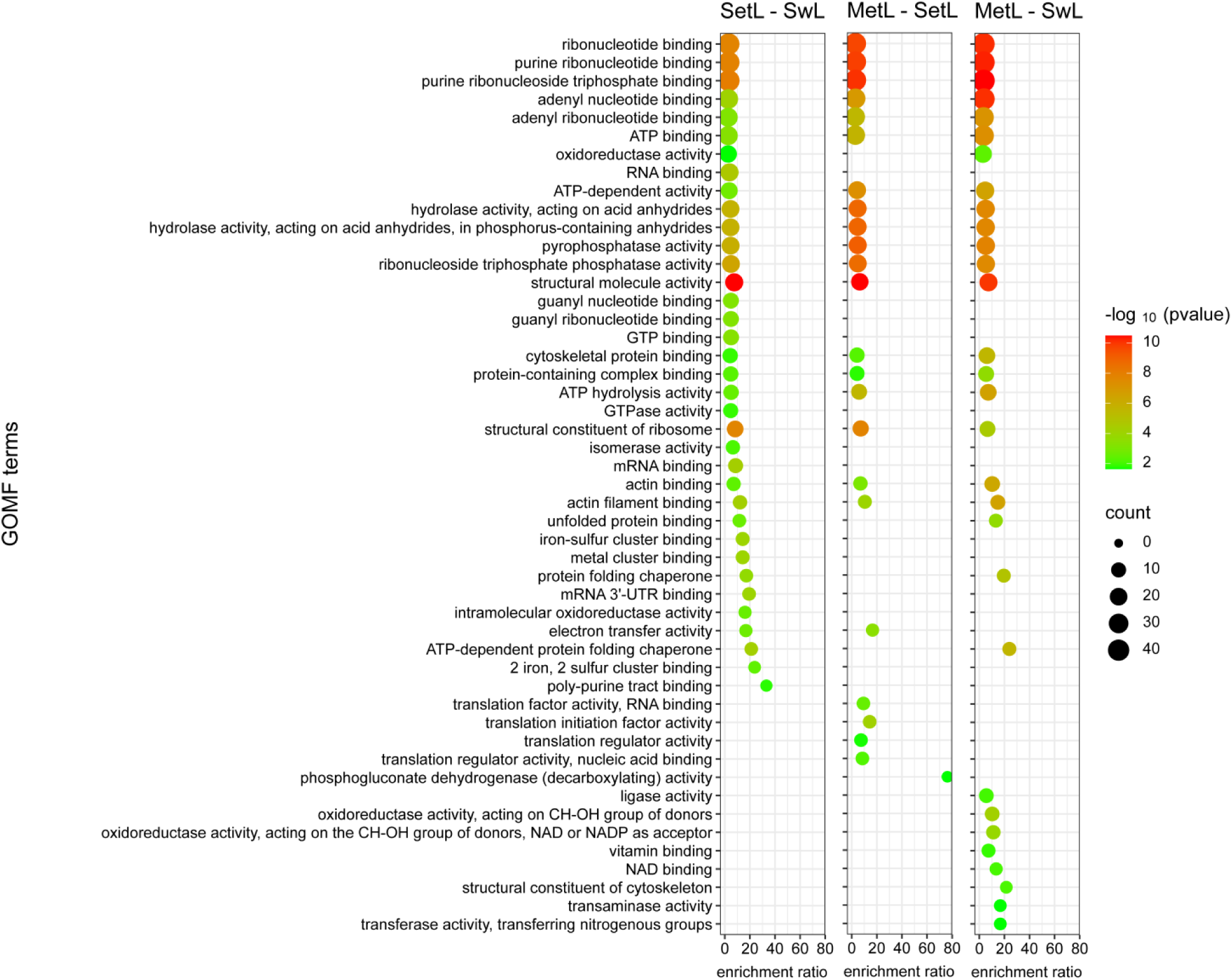
Gene Ontology Molecular Function (GOMF) terms enrichment analysis of differentially expressed proteins across the experimental comparisons. The dot plots show enriched GOMF terms for the comparisons SetL vs SwL, MetL vs SetL, and MetL vs SwL. The x-axis indicates the enrichment ratio, while the y-axis lists the enriched GOMF terms. The color gradient represents the statistical significance (−log₁₀ p-value), with red indicating higher significance, and the dot size corresponds to the number of proteins associated with each term.

**Figure S3.**
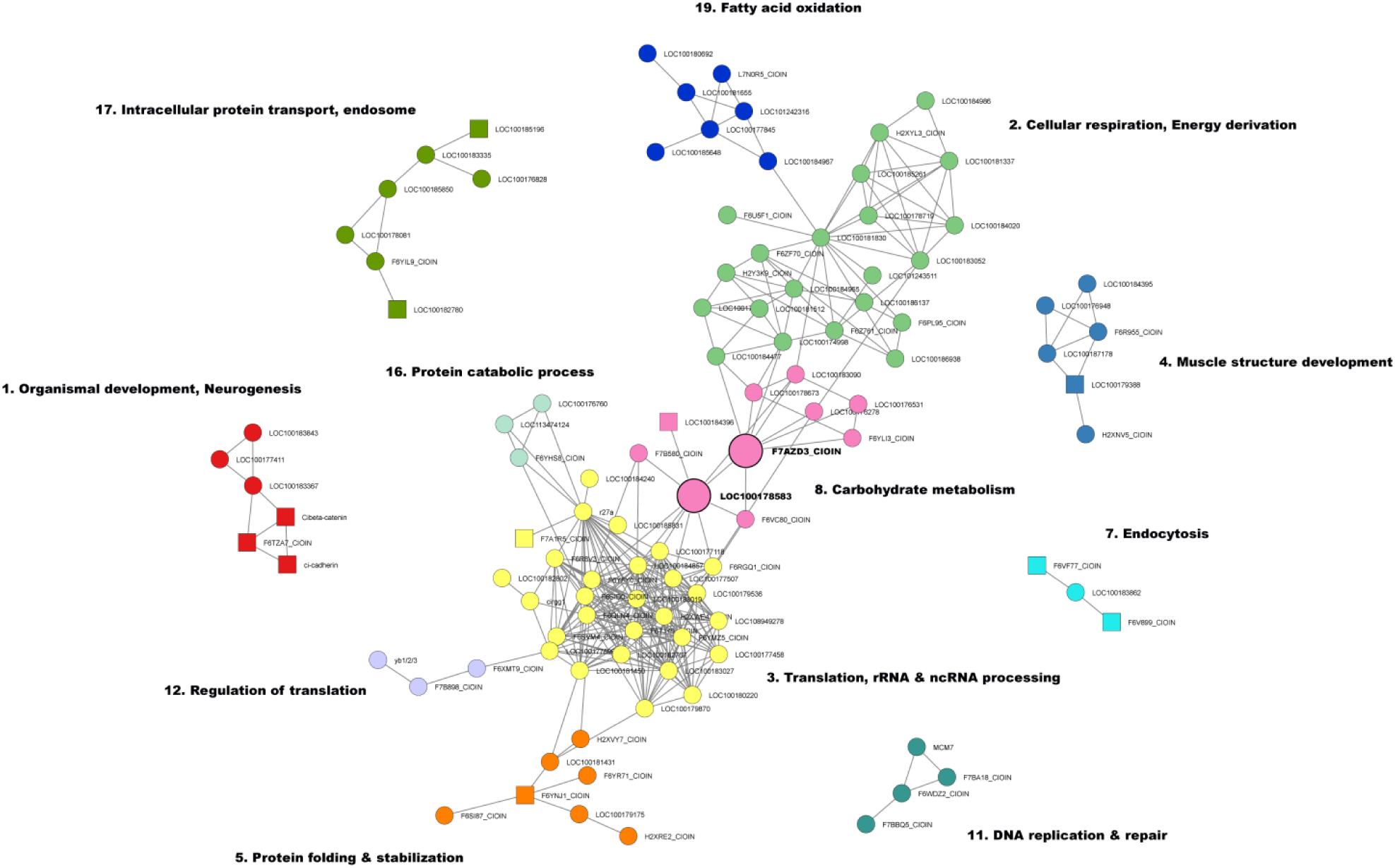
Network analysis of differentially expressed proteins in the MetL-SetL comparison. Node colors correspond to functional clusters (communities), and lines represent connections between proteins. Node sizes indicate the main connector hubs (see Table 6). Square nodes represent proteins annotated as “Neuronal” (see Suppl. Table S2). Major clusters are annotated with the most representative biological process names, highlighting key functional modules.

**Figure S4.**
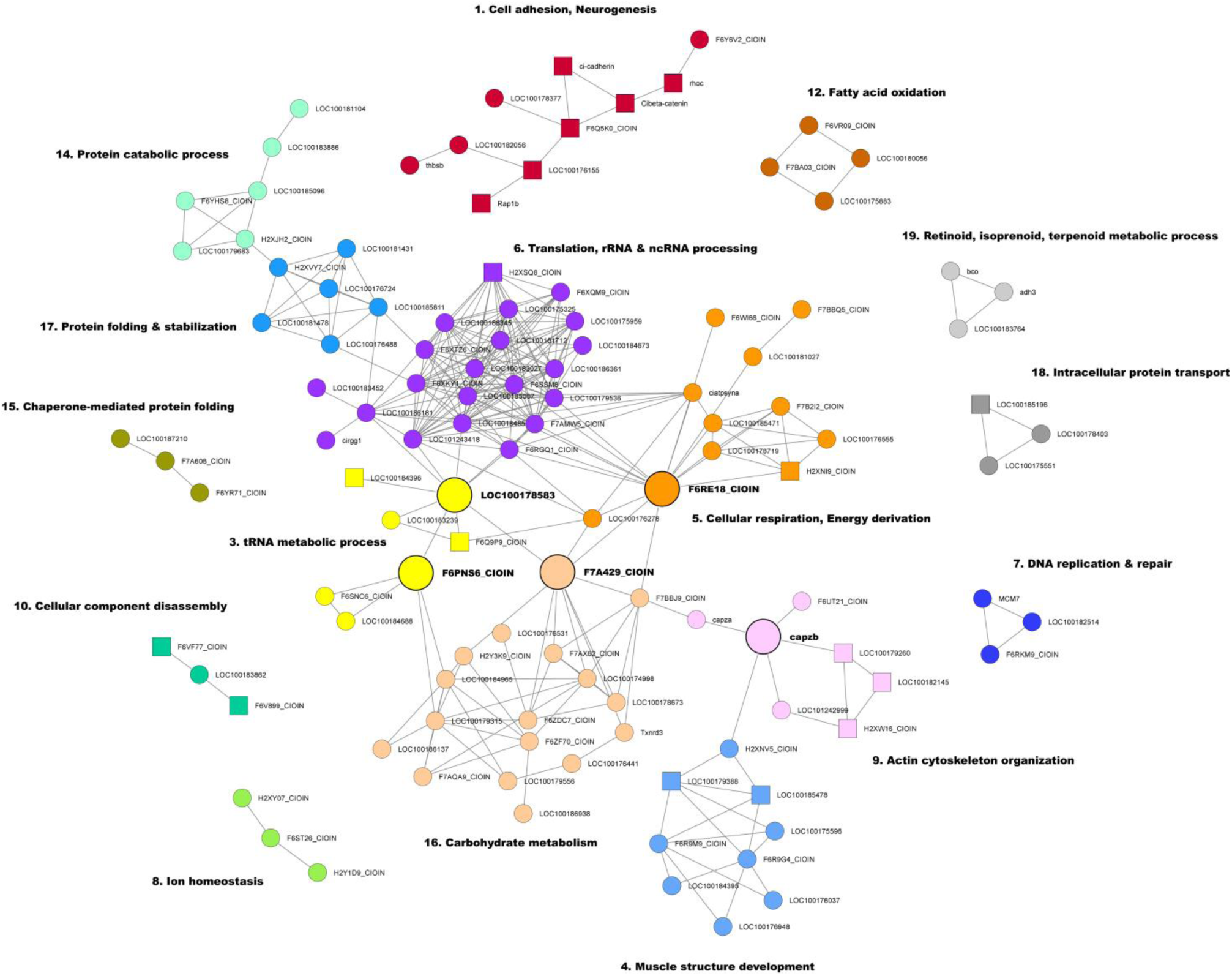
Network analysis of differentially expressed proteins in the MetL-SwL comparison. Node colors correspond to functional clusters (communities), and lines represent connections between proteins. Node sizes indicate the main connector hubs (see Table 6). Square nodes represent proteins annotated as “Neuronal” (see Suppl. Table S2). Major clusters are annotated with the most representative biological process names, highlighting key functional modules.

**Figure S5.**
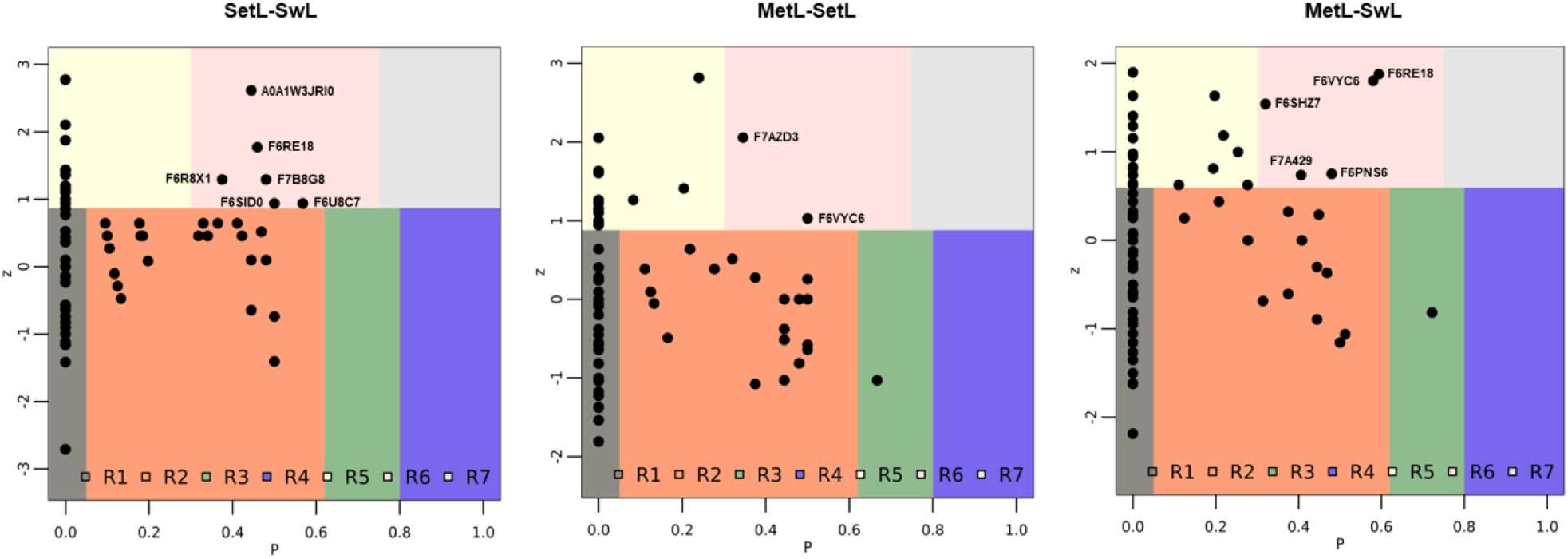
Functional cartography of the SetL-SwL, MetL-SetL and MetL-SwL networks. The maps show the distribution of proteins (nodes) within the network based on their participation coefficient (P, x-axis) and within-module degree z-score (z, y-axis). Nodes are categorized into seven distinct categories (R1 to R7) according to their functional connectivity profiles, each represented by a different color. This classification highlights the diversity of node roles in the network’s modular structure, distinguishing between nodes and hubs. The proteins classified as connector hubs (R6) in each network are labelled with the corresponding UniProtKB ID. Legend: R1 = Ultra-peripheral nodes; R2 = Peripheral nodes; R3 = Connector nodes; R4 = Kinless nodes; R5 = Provincial hubs; R6 = Connector hubs; R7 = Kinless hubs.

